# Volume loss and recovery in bovine knee meniscus loaded in circumferential tension

**DOI:** 10.1101/2023.02.24.529590

**Authors:** John M. Peloquin, Michael H. Santare, Dawn M. Elliott

**Affiliations:** Department of Biomedical Engineering, University of Delaware, Newark, Delaware; Department of Mechanical Engineering, University of Delaware, Newark, Delaware

## Abstract

Load-induced volume change is an important aspect of knee meniscus function because volume loss creates fluid pressure, which minimizes friction and helps support compressive loads. The knee meniscus is unusual amongst cartilaginous tissues in that it is loaded not only in axial compression, but also in circumferential tension between its tibial attachments. Despite the physiologic importance of the knee meniscus’ tensile properties, its volumetric strain in tension has never been directly measured, and predictions of volume strain in the scientific literature are inconsistent. In this study, we apply uniaxial tension to bovine knee meniscus and use biplanar imaging to directly observe the resulting 3D volume change and unloaded recovery, revealing that tension causes volumetric contraction. Compression is already known to also cause contraction; therefore, all major physiologic loads compress and pressurize the meniscus, inducing fluid outflow. Although passive unloaded recovery is often described as slow relative to loaded loss, here we show that at physiologic strains the volume recovery rate in the meniscus upon unloading is faster than the rate of volume loss. These measurements of volumetric strain are an important step towards a complete theory of knee meniscus fluid flow and load support.

## 1. Introduction

The knee meniscus likely depends on interstitial fluid pressure (“fluid load support”) to support compressive loads under physiologic conditions (Baro et al., 2012), similar to articular cartilage (Ateshian et al., 1994; Ateshian and Wang, 1995; Macirowski et al., 1994; Soltz and Ateshian, 2000; Soltz and Ateshian, 1998; Spilker et al., 1992; Zarek and Edwards, 1963). Positive interstitial fluid pressure reduces both the share of the load carried by the solid matrix and the tissue’s friction coefficient (Ateshian et al., 1994; Ateshian and Wang, 1995; Macirowski et al., 1994; McCutchen, 1959; Zarek and Edwards, 1963) which is believed to be important in preventing tissue damage (Ateshian et al., 1994; Macirowski et al., 1994; Mow et al., 1994; Soltz and Ateshian, 1998). However, with prolonged loading, fluid is lost and the pores gradually collapse, increasing strain and stress in the solid matrix (Ateshian et al., 1994; Mow et al., 1980; Zarek and Edwards, 1963). Significant and sustained fluid load support in the meniscus thus requires both volume loss and recovery: (1) when loaded, the meniscus must contract and exude fluid, decreasing in volume; and (2) when unloaded, it must recover the lost volume and fluid quickly enough to reset itself for the next loading cycle.

The change in meniscus volume during loading depends on both the applied loading and the mechanical properties of the porous solid matrix. Physiologic loading of the knee meniscus consists of both axial compression and circumferential tension (“hoop stress”) between its anterior and posterior attachments (coordinate system defined in Figure 1). Unconfined compression and indentation tests indicate that unconfined axial compression of the meniscus results in volume loss (Danso et al., 2018; Nguyen and Levenston, 2012; Sweigart et al., 2004). In vivo imaging and cadaver knee joint tests also indicate that cyclic gait loading causes the meniscus to lose volume overall (Benfield et al., 2022; Kessler et al., 2008, 2006). The specific effect of circumferential tension on meniscus volume, however, is less clear and is the main subject of this paper.

**Figure 1:**
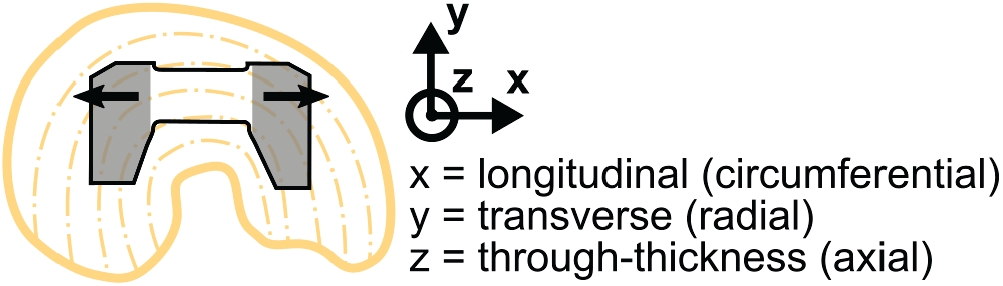
Schematic of circumferential uniaxial tension specimens, showing where tissue specimens were cut from the donor menisci, and the coordinate system used throughout this paper. Dashed lines = prevailing local circumferential fiber direction, gray shading = gripped region, and arrows = tension direction.

Constitutive modeling of the meniscus varies a great deal in its treatment of how tension translates into volume strain. Isotropic elasticity requires by definition that tension cause a volume increase, whereas material anisotropy removes this restriction. Even though knee meniscus tissue is anisotropic (Goertzen et al., 1997; LeRoux and Setton, 2002; Peloquin et al., 2018, 2016; Proctor et al., 1989; Tissakht and Ahmed, 1995), many computational works treat the meniscus as isotropic and therefore implicitly assume that tension increases volume. Even among those works that use an anisotropic model, many set the Poisson ratio parameters < 0.5 (Gu and Li, 2011; Haut Donahue et al., 2003; Vadher et al., 2006; Vaziri et al., 2008; Wilson et al., 2003; Yao et al., 2006; Zielinska and Donahue, 2006), with the same result. Only a few works use constitutive models that predict a volume decrease in circumferential tension (LeRoux and Setton, 2002; Spilker et al., 1992; Upton et al., 2006).

The available experimental evidence related to tension-induced volume change is not definitive. Volume is a 3D measurement and cannot be determined from 2D data without assuming a relationship, such as transverse isotropy, that prescribes the behavior of the unobserved dimension. Prior work with the meniscus measured planar (2D) deformations from three separate uniaxial tension tests, each in a different direction (Goertzen et al., 1997; LeRoux and Setton, 2002). These results suggest that circumferential tension would decrease meniscus volume. Unfortunately, reported material parameters differ substantially between the two studies and, as noted therein, when the values from each test are combined they do not satisfy the symmetry and positive definiteness requirements for a transversely isotropic linear elastic material (Goertzen et al., 1997; LeRoux and Setton, 2002). The meniscus’ volume change and general 3D deformation in tension have not yet been observed directly and remain uncertain. To summarize the state of the literature, simulation studies of the meniscus tend to assume that tension *increases* meniscus volume, the best available experimental data suggests with a great deal of uncertainty that tension instead *decreases* volume, and there is a clear need to resolve the conflict through direct observation of the actual tension-induced volume change.

The rate and degree of volume recovery on unloading, with accompanying re-imbibition of exuded fluid, is just as important as volume change under load. Knee loading is cyclic, with short periods of unloading during the gait cycle and longer periods of unloading during rest. Sustained fluid pressurization depends on volumetric and thus fluid recovery, priming the tissue for the next loading cycle. Human meniscus volume decreases during a 20 km run (Kessler et al., 2008, 2006), so recovery within the gait cycle itself is not always complete and unloaded recovery during rest is probably physiologically important. Prior reports of unloaded recovery of meniscus are very limited, consisting of one representative indentation creep recovery curve for rabbit meniscus (Sweigart et al., 2004; Sweigart and Athanasiou, 2005a) and the aforementioned human MRI study, which showed recovery of ∼ 50% of lost meniscus volume with 60 min bed rest post-run (Kessler et al., 2008). Unloaded recovery of tendon, spine, and articular cartilage has received considerably more attention (Amin et al., 2016; Bezci et al., 2020; Bezci and O‘Connell, 2018; Crisco et al., 1997; Duenwald et al., 2009; Ekström et al., 1996; Graf et al., 1994; Han et al., 2000; Johannessen et al., 2004; Malko et al., 2002; McGill and Brown, 1992; O‘Connell et al., 2011; Schmidt et al., 2016; Tyrrell et al., 1985), although generally without direct measurement of 3D deformation and thus volumetric recovery. Considering all the above, the objectives of this work were (1) to measure 3D strain and volume change in the meniscus over time in circumferential tension and after unloading and (2) to compare the rate of volume change under tension to the rate of unloaded volume recovery.

## 2. Methods

### 2.1. Specimen characteristics

Ten bovine menisci (5 medial, 5 lateral, age 3 months to 12 years), were acquired from 3 different commercial sources (Animal Technologies, Tyler, TX; Herman’s Quality Meats, Newark, DE; and Green Village Packing Co., Green Village, NJ) to increase sample representativeness. Menisci were stored in double-bagged vacuum-sealed pouches at −20 °C until use. On the day a meniscus was tested, it was sliced into ∼ 3 mm thick slices in the axial plane using a deli slicer (Creechley et al., 2017; Henderson et al., 2022; Wale et al., 2021), and a modified dogbone specimen was cut from the mid-axial slice (Figure 1) (Peloquin et al., 2018, 2016). The elongated gripped region on the inner side of the specimen allowed more of the innermost circumferential fibers’ length to be gripped; this has been shown to increase measured tensile modulus and strength compared to transversely symmetric dogbone-shaped specimens (Peloquin et al., 2016).

Immediately after the specimen was cut, mean specimen thickness was measured using a scanning laser interferometer (Favata, 2006; Szczesny et al., 2012) with 5 equally spaced passes over the gauge region. The other dimensions were measured with digital calipers. Across all specimens, gripto-grip length = 14.7–16.9 mm (16.0 ± 0.9 mm), length of the parallel-sided gauge region = 10.4– 11.2 mm (10.7 ± 0.3 mm), central width = 5.3–10.1 mm (6.7 ± 1.6 mm), thickness = 1.7–4 mm (2.9 ± 0.6 mm), and central cross-sectional area = 11.8–27.6 mm^2^ (19.1 ± 5.2 mm^2^). Overall specimen length was ∼ 38 mm. The specimen-specific cross-sectional area from these measurements was later used to calculate circumferential stress, σ_xx_.

### 2.2. Test setup

Test setup began with mounting the specimen in tension grips on the bench. Two sheets of 400 grit cloth sandpaper were placed between the specimen and the serrated grip faces. The grips were connected by rigid plates that protected the specimen from accidental load during setup. To maintain adequate clamping pressure throughout the test, the grips were tightened three times in 10 minute intervals (Peloquin et al., 2016; Schechtman and Bader, 1997; Swank et al., 2014). During this ∼ 25 minute period the specimen was wrapped in gauze dampened with phosphate buffered saline (PBS). The gripped specimen was transferred to an Instron 5943 and immersed in a bath filled with commercially prepared PBS (Fisher BioReagents BP3994). All fixture connections (which were the socket, pin, and lock nut type) were tightened with 1 kN tension across the rigid plates connecting the grips to reduce load string laxity (load string stiffness = 3.5 kN/mm). The rigid plates were then removed. One camera (Basler a 102f CCD; Navitar Zoom 7000) was then focused on the front face of the specimen and a second (Canon EOS Rebel T7i DSLR; Canon EF 180mm f/3.5L) on the side of the specimen, completing setup of the apparatus (Figure 2).

**Figure 2:**
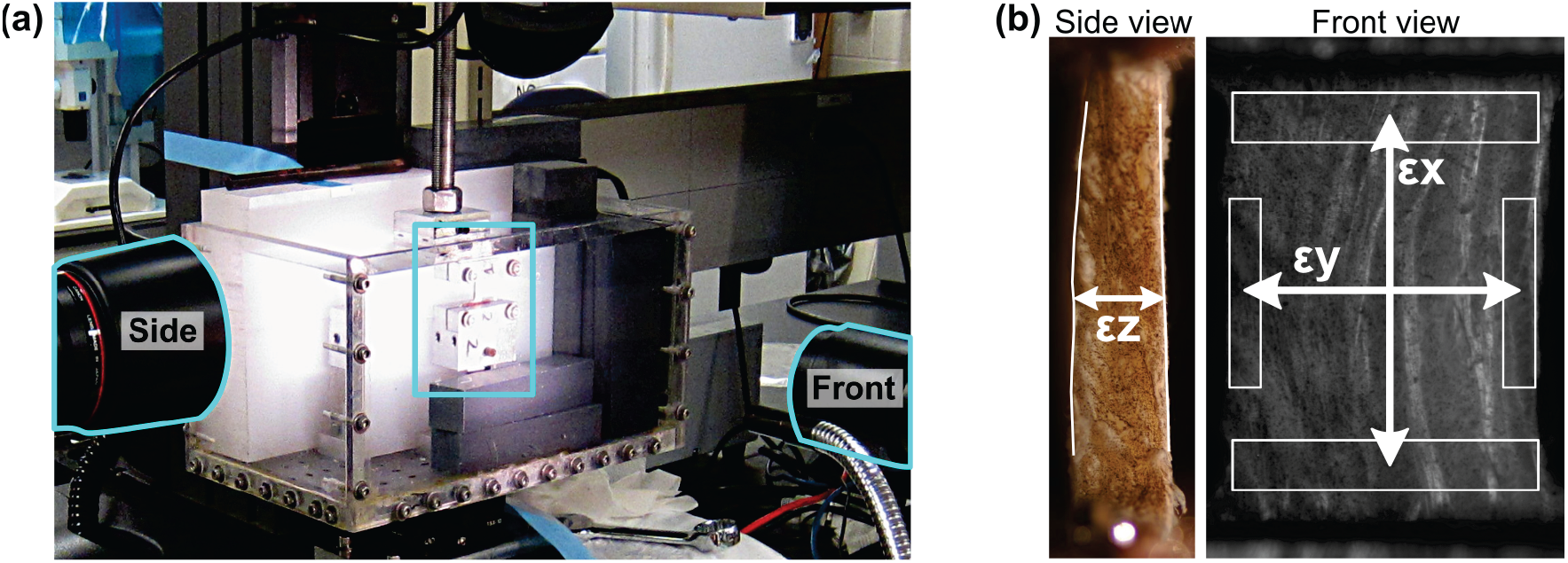
Uniaxial tension test setup. (a) Front and side camera positions. Cameras are outlined in cyan and the grips & specimen are marked with a cyan box. (b) Strain components were measured from biplanar video of the specimen’s deformation using mean distance between manually labeled edges (ε_z_) or distance between the centroids of regions of interest (ROIs) tracked by image registration (ε_x_ and ε_z_).

Strain was set to zero under a 0.5 N tensile preload. In tensile tests of soft tissue it is typical for the tissue strain to be less than the grip-to-grip strain (“grip strain”; crosshead displacement / between-grip distance at preload) (Firminger and Edwards, 2021; Han et al., 2016, 2013; Lee et al., 2017; Peloquin et al., 2018, 2016; Szczesny and Elliott, 2014). In the present work, the discrepancy between tissue and grip strain was not caused by grip slip in the usual sense, but rather by the effective grip line being somewhere within the gripped region rather than exactly at the grip face. In other words, the effective gauge length was greater than the nominal gauge length. Therefore, for each specimen, the crosshead displacement was calibrated incrementally over 3 tension–unloading cycles to produce the desired longitudinal tissue strain. After each calibration cycle, the tissue strain produced by the applied crosshead displacement was measured using digital image correlation (Vic-2D’s extensometer tool, 0.7 mm correlation window; Correlated Solutions, Columbia, SC), and a new crosshead displacement was calculated for the next cycle according to the observed tissue strain:grip strain ratio. The final tissue strain:grip strain ratio thus calculated (ranging from 0.48 to 0.91) was carried forward to the tensile test protocol, providing more accurate control of tissue strain. Note that this crosshead calibration was only used for actuator control; for analysis, the actual tissue strain was directly measured during post-processing of the data (see below).

### 2.3. Tensile test protocol

The tensile test consisted of four phases of uniaxial tension in the circumferential (x) direction (Figure 1), each phase consisting of either stress relaxation or creep loading followed by unloaded recovery (Figure 3). The first stress relaxation phase was for preconditioning at the full target strain level. The middle (“main”) stress relaxation and creep phases were the main subject of analysis. The final stress relaxation verification phase was used to verify that the specimen did not accrue damage during the creep phase (Johannessen et al., 2004; Koeller et al., 1984; Showalter et al., 2014), and indeed there was negligible change in stress-bearing capacity between the main stress relaxation and the verification phases (Supplemental Figure 3).

**Figure 3:**
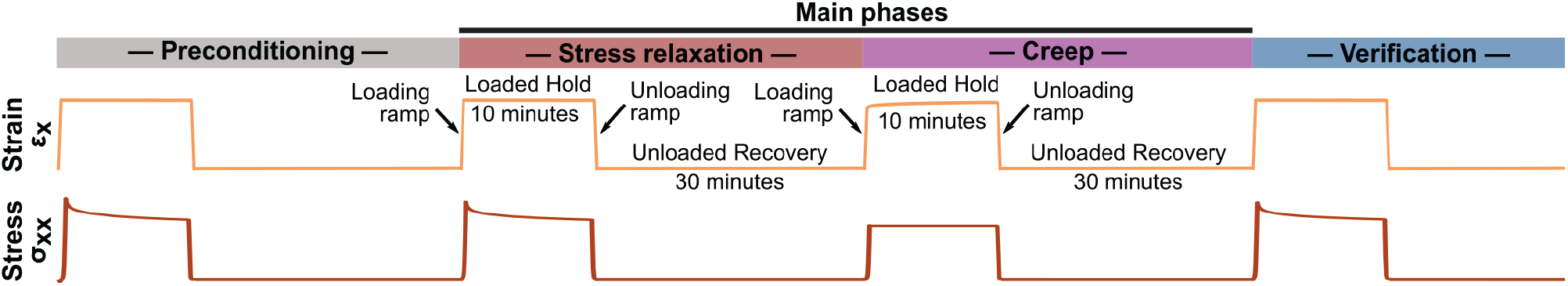
Uniaxial circumferential tension test protocol schematic. The preconditioning and verification phases are identical to the main stress relaxation phase.

During each stress relaxation phase, a crosshead displacement was applied to produce a tissue longitudinal engineering strain ε_x_ = 6.4%, according to the tissue strain:grip strain ratio observed in that specimen’s calibration cycles, and held for 10 minutes. During the creep phase, a crosshead displacement equal to 7/8 of that applied in the stress relaxation phases was applied and the resulting load was held for 10 minutes. Loading and unloading ramps were applied at crosshead displacement rate = 0.1 mm/s. The stress relaxation target strain of 6.4% was chosen for consistency with similar prior work (LeRoux and Setton, 2002), and the creep load level was chosen based on preliminary testing to avoid the tissue strain growing to exceed the preconditioned strain level, as excess strain might change the properties of the tissue mid-test by creating new damage or plastic deformation. The most representative set of time series data is shown in Figure 4.

**Figure 4:**
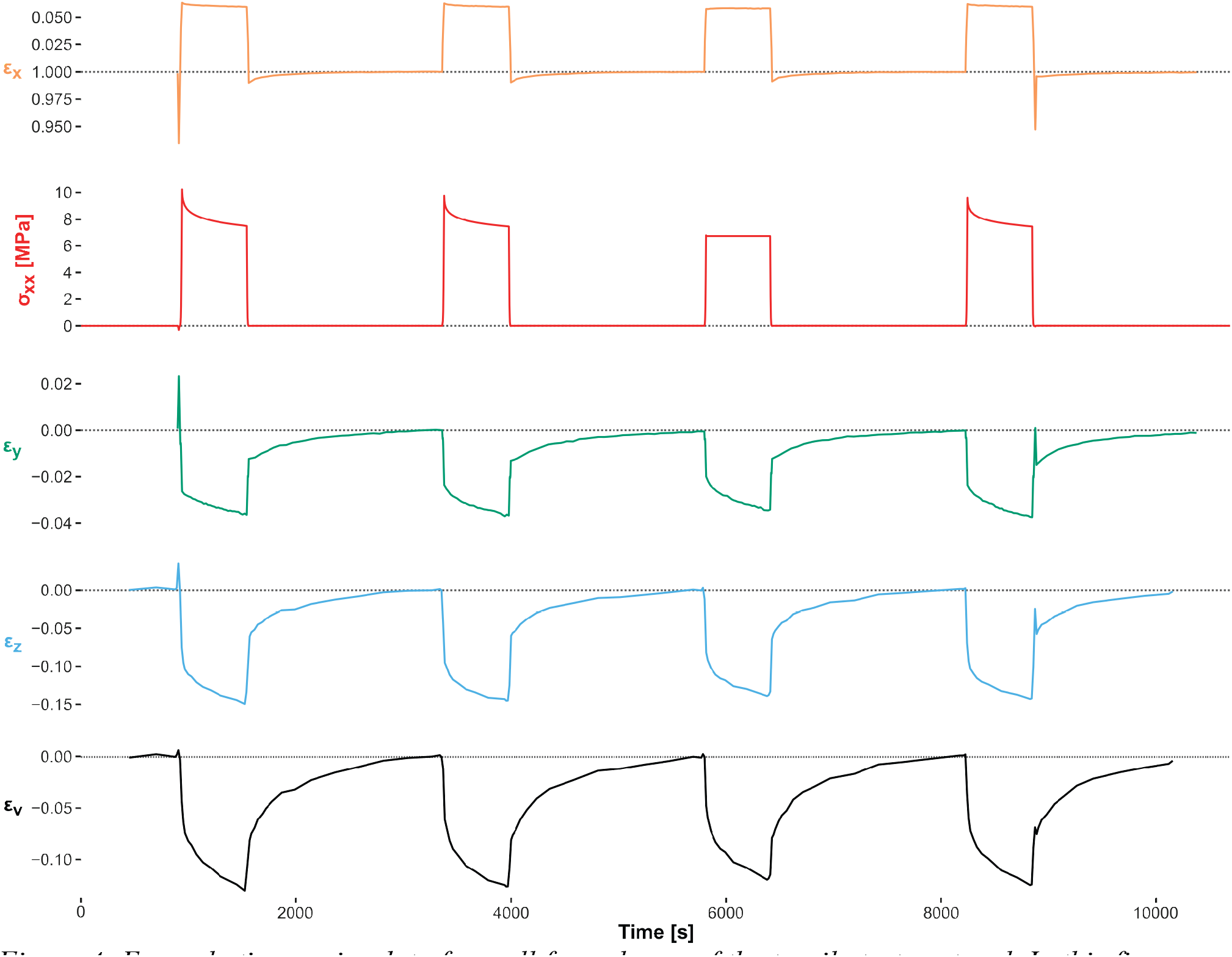
Example time series data from all four phases of the tensile test protocol. In this figure, the end of preconditioning was used as the reference configuration for strain. Time series plots for all specimens are available in this paper’s supplemental information.

### 2.4. 3D strain measurement

To calculate volume change, the engineering strains ε_x_, ε_y_, and ε_z_ were measured from video of the front and side of the specimen (Figure 2). ε_x_ and ε_y_ were measured from the displacement of the ROIs shown in Figure 2b. The ROIs were manually drawn in the reference configuration’s image (Figure 2b) and tracked through all other images by affine registration of the entire ROI using Advanced Normalization Tools (ANTs) (Avants et al., 2014, 2011). Each x-strain ROI spanned the specimen width and 10% of the grip-to-grip length. Each y-strain ROI spanned 40% of the grip-to-grip length and 10% of the specimen width. ANTs was wrapped in a Python library (https://github.com/jpeloquin/postmech) that initialized each registration with the values obtained for the immediately preceding image, enabling the registration to follow large deformations. This method was developed because Vic-2D exhibited extensive decorrelation errors and tracking loss when applied to the full image dataset, such that selection bias (discarding harder-to-track regions, e.g., high-strain regions) was a major concern. ε_z_ was measured from the mean displacement of the specimen edges in the side video of the test, which were labeled by hand (Figure 2b). Volume “strain” ε_v_ was calculated as ε_v_ = *J* − 1, where *J* is the volume ratio (in general, the determinant of the deformation tensor; in this specific test, the product of the stretch ratios λ_x_, λ_y_, and λ_z_).

### 2.5. Statistics

To analyze how strain and volume (ε_x_, ε_y_, ε_z_, and ε_v_) changed as circumferential tension was applied and removed, strain during the two main stress relaxation and creep phases was re-referenced, by division of the stretch ratios, to the initial state immediately preceding the phase’s loading ramp.

With this reference configuration, subsequently referred to as the “phase reference configuration”, ε = 0 means no change relative to the start of the phase. All tests of whether strain changed at all within a test phase, and in which direction, were therefore done by comparison to zero with the one-sample t-test. The rate of strain change during each loaded hold and unloaded recovery hold was quantified using the slope of a linear fit to ε vs. time during the first and last 100 s of the hold (Voinier et al., 2022), as 100 s was the shortest time span that guaranteed representative sampling of the DSLR time lapse. For comparison with historical results, the time to achieve 63% of the hold’s total change was also calculated (“characteristic time”; Supplemental Figures 1 and 2).

All comparisons of values between time points not involving the reference configuration were done by repeated measures ANOVA followed by paired t-tests when overall significance was indicated, including load type (stress relaxation or creep) as a factor. Statistical tests were done in R 4.2.1 and checked in JMP Pro 16 (Müller and Wickham, 2022; R Development Core Team, 2022; Wickham et al., 2022; Wickham, 2016). According to recent recommendations, significance was set at p < 0.005 and tendency at 0.005 ≤ p < 0.05 (Benjamin et al., 2017; Ioannidis, 2005). Unless otherwise indicated, values are reported as mean ± s.d. pooled across stress relaxation and creep, as they often produced indistinguishable y- and z-strain distributions.

## 3. Results

Volume tended to decrease with application of uniaxial tension during the loading ramps. At the end of the loading ramps (i.e., start of the loaded hold), for both stress relaxation and creep, ε_v_ = −0.027 ± 0.028 (≠ 0, p < 0.05) (Figure 5, start). Only two specimens increased in volume during at least one loading ramp, and this increase was temporary. During the loaded holds (Figure 5), volume decreased for every specimen (start to end Δε_v_ = −0.048 ± 0.030; ≠ 0, p < 0.005). At the end of the loaded holds, this volume decrease was large enough that every specimen showed volume loss relative to the phase reference configuration (end ε_v_ = −0.084 ± 0.047; ≠ 0, p < 0.005).

**Figure 5:**
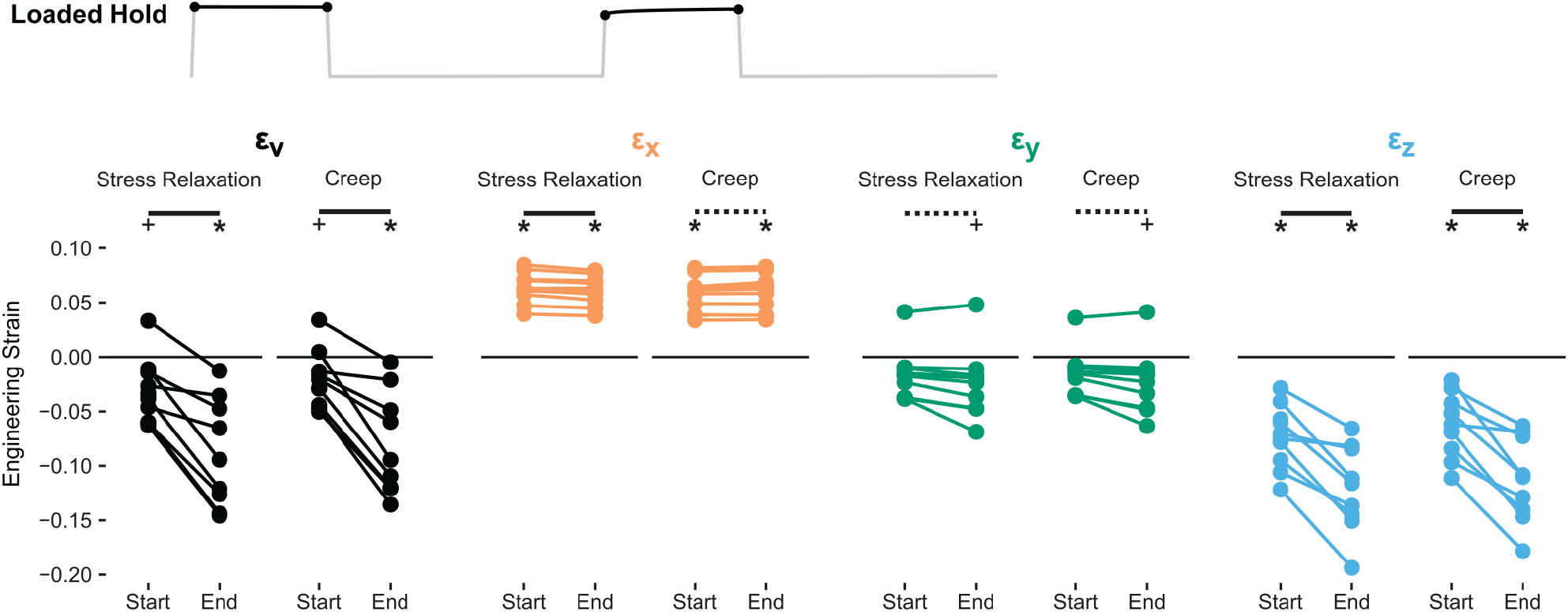
Engineering strain in the gauge region at the start and end of each loaded hold (10 minutes duration). Test for mean ε ≠ 0: +, p < 0.05; *, p < 0.005. Test for mean start to end Δε ≠ 0: dashed overbar, p < 0.05; solid overbar, p < 0.005.

The main cause of volume loss in tension was through-thickness contraction in ε_z_ (Figure 5). ε_z_ consistently decreased by a large amount during both the loading ramp (loaded hold start ε_z_ = −0.067 ± 0.029; ≠ 0, p < 0.005) and the loaded hold (loaded hold Δε_z_ = −0.048 ± 0.025; ≠ 0, p < 0.005). ε_y_ also tended to contribute to volume loss but to a much lesser degree than ε_z_ (loaded hold Δε_y_ = −0.008 ± 0.009; ≠ 0, p < 0.05).

The unloading ramp produced a small amount of volume recovery in all but one specimen (Figure 6, start to end), but the mean change was not significantly different from zero. The unloading ramp drove ε_x_ to slightly less than zero, and also caused ε_y_ to become not significantly different from zero (Figure 6, end). ε_z_ reversed about half of its contraction during the unloading ramp but remained substantially less than zero at the end of the ramp (end ε_z_ = −0.057 ± 0.035; ε_z_ ≠ 0, p < 0.005).

**Figure 6:**
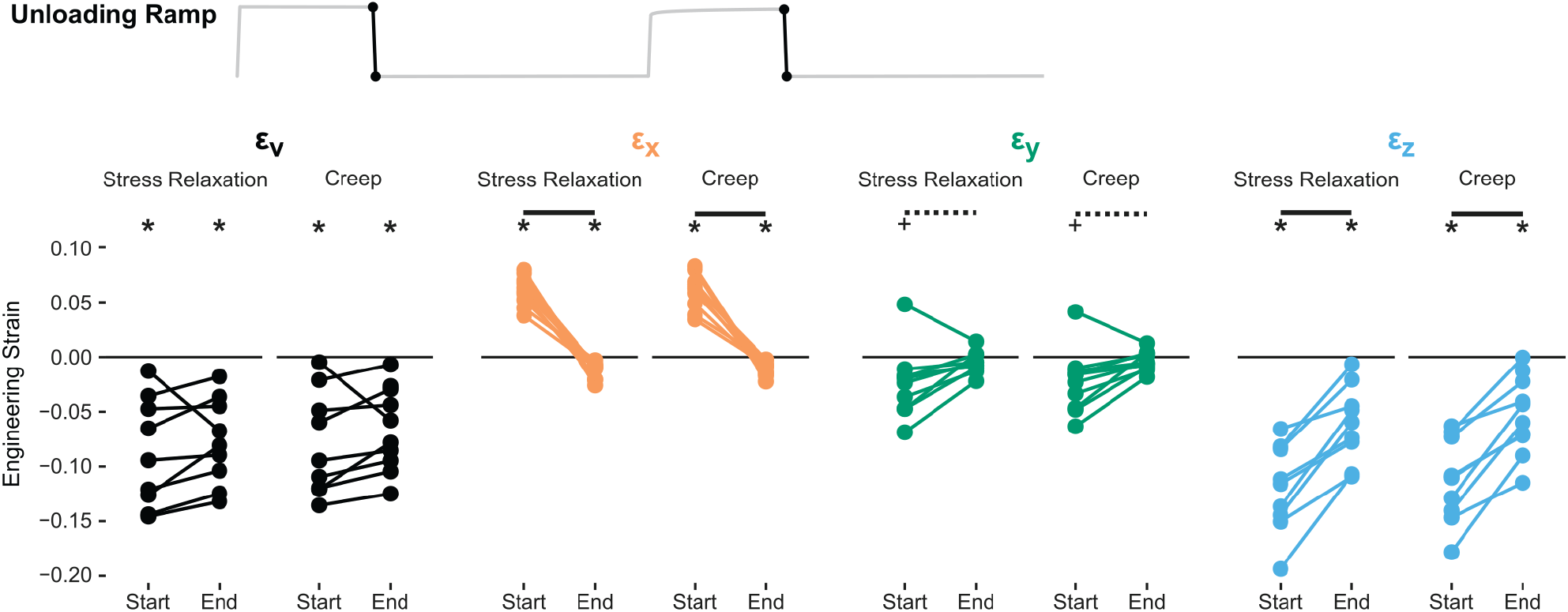
Engineering strain in the gauge region at the start and end of each unloading ramp. Test for mean ε ≠ 0: +, p < 0.05; *, p < 0.005. Test for mean start to end Δε ≠ 0: dashed overbar, p < 0.05; solid overbar, p < 0.005.

Overall, volume changed little during the unloading ramp because reversal of existing expansion in ε_x_ was balanced by reversal of existing contraction in ε_y_ and ε_z_.

During unloaded recovery (Figure 7), mean ε_v_ across specimens became nearly zero after stress relaxation, albeit with considerable variability (end ε_v_ = 0.002 ± 0.013) and nearly but not exactly zero after creep (end ε_v_ = 0.004 ± 0.005; ≠ 0, p < 0.05). ε_x_ reversed its slight residual contraction and became almost exactly zero (end ε_x_ = 0.003 ± 0.009, across both stress relaxation and creep). ε_y_ remained not significantly different from zero, although its distribution narrowed somewhat (end ε_y_ = 0.001 ± 0.005, stress relaxation and creep). ε_z_ reversed its existing contraction, becoming not significantly different from zero after stress relaxation (end ε_z_ = 0.002 ± 0.006) and nearly zero after creep (end ε_z_ = 0.003 ± 0.003; ≠ 0, p < 0.05). ε_z_ was thus the main cause of volume recovery during unloaded recovery, in the same way that it was the main cause of volume loss during the loaded holds.

**Figure 7:**
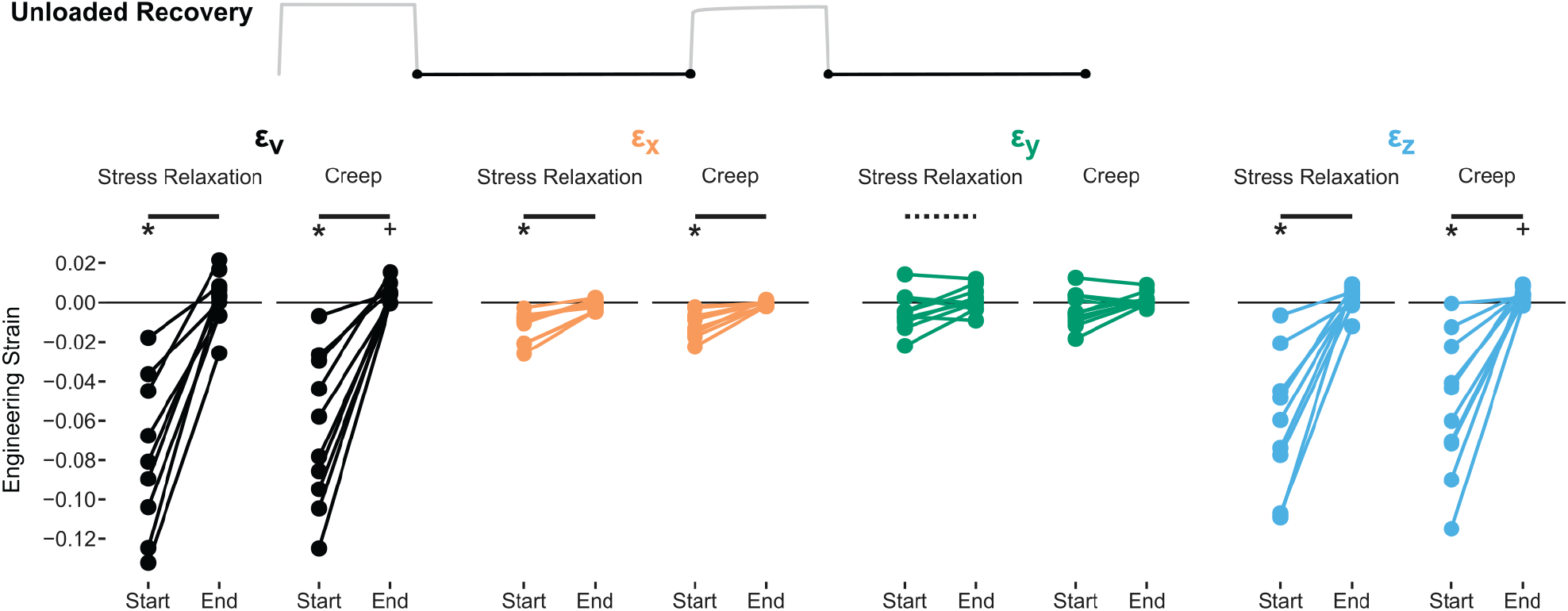
Engineering strain in the gauge region at the start and end of each unloaded recovery (30 minutes duration). Test for mean ε ≠ 0: +, p < 0.05; *, p < 0.005. Test for mean start to end Δε ≠ 0: dashed overbar, p < 0.05; solid overbar, p < 0.005.

The rate of change in ε_v_ and ε_z_ was rapid during the start of each loaded hold as well as during the start of each unloaded recovery (Figure 8). The rate of volume loss at the start of the loaded hold tended to be slightly greater than its rate of recovery at the start of unloaded recovery (p = 0.01). However, the rate of volume recovery during the start of each unloaded recovery greatly exceeded the rate of volume loss during the end of each loaded hold (p < 0.005). ε_z_, as the main contributor to ε_v_, followed the same pattern. ε_x_ was directly driven throughout much of the test, so its rate of change is not particularly interesting. The rate of change for ε_y_ was very small at all times.

**Figure 8:**
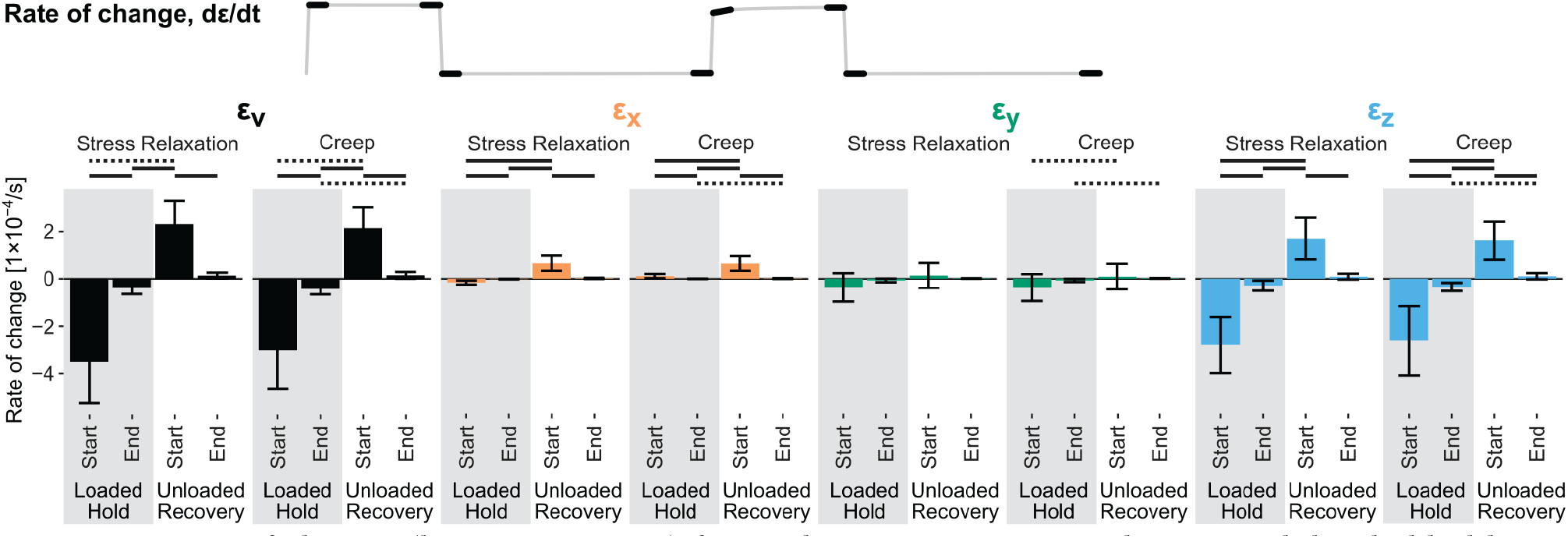
Rate of change (loss or recovery) for each strain component during each loaded hold or unloaded recovery. The rate of change was measured during the first and last 100 s of each hold as the slope of a least squares line fit. Test for mean loss ≠ mean recovery: dashed overbar, p < 0.05; solid overbar, p < 0.005. The rate of change during the end of unloaded recovery, which is at equilibrium, was not compared with rate of change during the start of the loaded hold.

Swelling over the main part of the test was measured as the change in each strain component from the end of the preconditioning phase to the end of the creep unloaded recovery, using the end of preconditioning as the reference configuration. On average, ε_v_ and ε_y_ did not significantly change over the course of the test (Figure 9; Δε_v_ = 0.006 ± 0.017; Δε_y_ = 0.003 ± 0.010). ε_x_ tended to decrease slightly (Δε_x_ = −0.002 ± 0.003; Δε_x_ ≠ 0, p = 0.05) and ε_z_ tended to increase slightly (Δε_z_ = 0.006 ± 0.005; Δε_z_ ≠ 0, p = 0.01). Although the average changes were small, there was significant variation between individual specimens, with some shrinking and others expanding over the course of the test.

**Figure 9:**
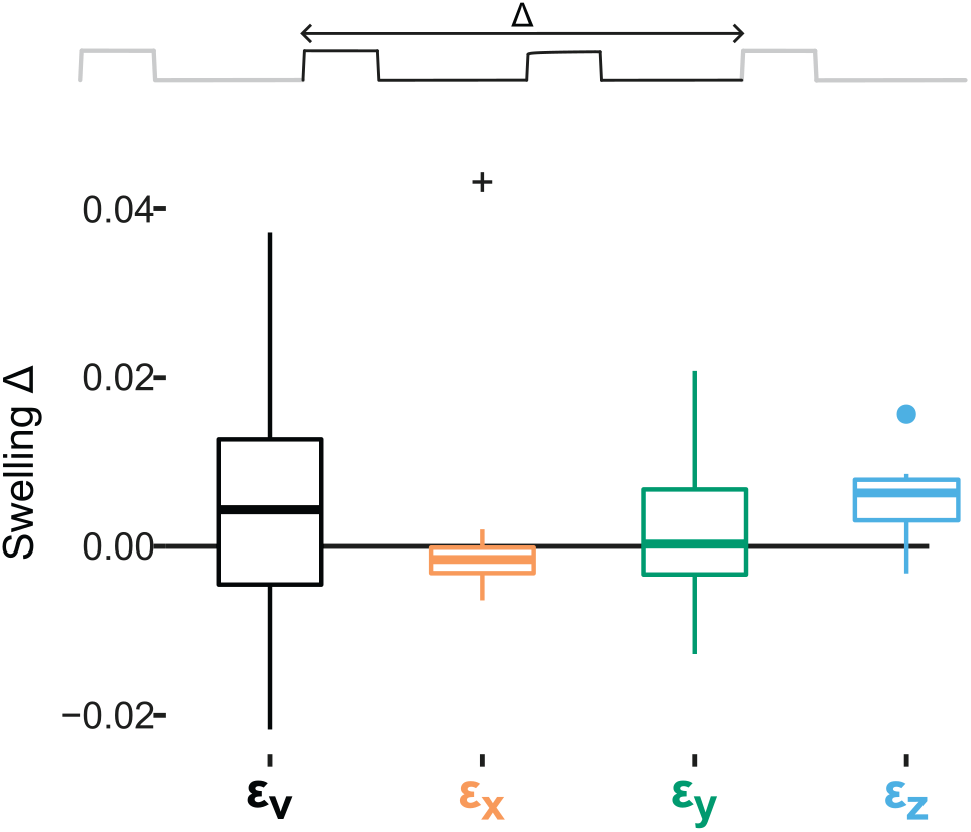
Change in strain over the main test phases, measured from the end of preconditioning to the end of unloaded recovery from creep. In this figure, the end of preconditioning is used as the reference configuration. Box plot line = median, hinges = quartiles, whiskers = min/max of values nearer than 1.5 IQR from the median, and points = values further than 1.5 IQR from the median. Test for mean ε ≠ 0: +, p < 0.05; *, p < 0.005.

## 4. Discussion

### 4.1. Overview

Direct measurement of volume strain in meniscus tissue in uniaxial tension demonstrated that circumferential tension decreases tissue volume. Holding the load for 10 minutes approximately doubled the loss of volume. Volumetric contraction was primarily caused by contraction in the through-thickness direction (z), not the transverse direction (y). During unloaded recovery, the rate of volume recovery was rapid, faster than the rate of volume loss towards the end of each loaded hold but not quite as fast as the rate of volume loss at the very beginning of each loaded hold.

Volume loss in tension implies fluid pressurization and exudation, so these results indicate that circumferential tension in the meniscus is partially supported by interstitial fluid pressure.

### 4.2. Volume strain and strain components

Simultaneous measurement of all three normal strain components (ε_x_, ε_y_, and ε_z_) proved essential to obtain an accurate measurement of volume strain. Considering tensile tests of the meniscus and other fibrous soft tissues, almost all prior work reports planar (2D) measurements, often expressed as longitudinal–transverse Poisson ratios. 3D strain has been reported twice, for articular cartilage subjected to uniaxial tension, revealing different transverse (ε_y_) and through-thickness (ε_z_) strains (Chang et al., 1999; Woo et al., 1979). The present work found that in the meniscus, contraction in ε_z_ is similarly much greater than contraction in ε_y_. Therefore, although the meniscus is typically considered to be transversely isotropic (Freutel et al., 2015; Goertzen et al., 1997; Haut Donahue et al., 2003; LeRoux and Setton, 2002; Meakin et al., 2003; Mononen et al., 2015, 2013; Spilker et al., 1992; Upton et al., 2006; Vadher et al., 2006; Wheatley et al., 2015; Yao et al., 2006; Zielinska and Donahue, 2006), calculation of ε_v_ from x–y planar data (i.e., assuming ε_z_ = ε_y_, with transverse isotropy as the justification) will, for many specimens, produce an incorrect prediction of volumetric *dilatation* in tension rather than contraction (Figure 10).

**Figure 10:**
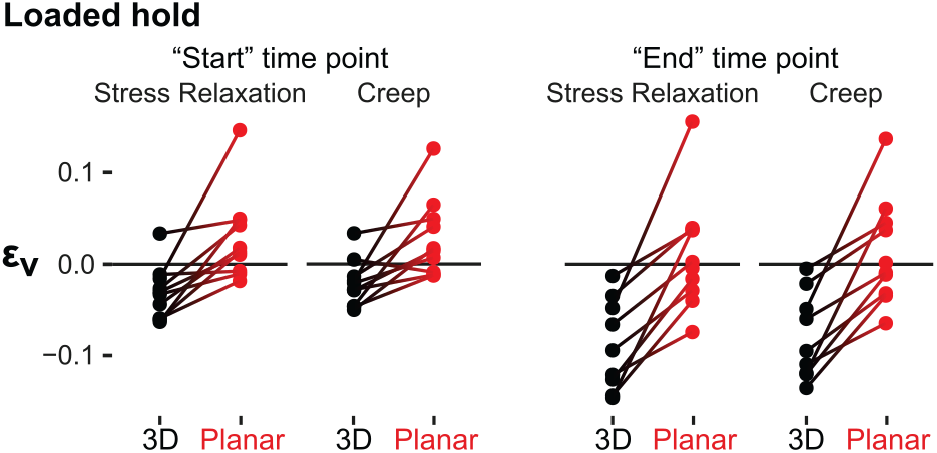
Comparison, for each specimen, of volume strain ε_v_ measured directly from ε_x_, ε_y_, and ε_z_ (“3D” ε_v_) to the prior approach of estimating ε_v_ from ε_x_ and ε_y_ alone under the assumption that the deformation is transversely isotropic (“planar” ε_v_).

The observed differences between ε_y_ and ε_z_ are likely influenced by grip effects, so these results only indicate that it is imprudent to assume transverse isotropy in an experimental context. The underlying constitutive law for meniscus tissue could still be transversely isotropic. Clamp-style grips induce through-thickness compression and, more importantly, restrict lateral deformation even in the midsubstance (Carniel et al., 2019). Even though tendon is supposed to be transversely isotropic, uniaxial tension tests of porcine deep flexor tendon produce lateral expansion and through-thickness contraction (Carniel et al., 2019). High fiber stiffness combined with low between-fiber shear stiffness causes grip-induced stress field perturbations in fibrous tissue to extend into the midsubstance unless the specimen aspect ratio is very large, on the order of 40:1 (Horgan and Simmonds, 1994; Reese et al., 2013; Skaggs et al., 1994b; Sun et al., 2005; Szczesny et al., 2015). In a pure uniaxial tension test, the transverse and through-thickness axes are completely unrestrained, but in real tissue tests this is not true. In the present test, the wider transverse direction is more constrained than the thinner through-thickness direction, so ε_y_ and thus ε_v_ probably contracted less than in a hypothetical pure uniaxial tension test.

Biological specimens are restricted in size and shape by the dimensions of the donor organ, which forces a uniaxial test to be non-ideal. Meniscus fiber bundles (fascicles) are 50–500 μm in diameter and, although generally circumferentially aligned, are dispersed in orientation (Andrews et al., 2013; Kelly et al., 1990; Petersen and Tillmann, 1998; Sweigart and Athanasiou, 2005a). A high aspect ratio can only be achieved by making the specimen very narrow, ∼ 1 mm wide, which reduces fiber continuity and makes it implausible to consider the material as a continuum (in the sense used in continuum mechanics) (Goertzen et al., 1997; Wale et al., 2021). Fiber continuity has also been directly shown to impact ultimate tensile stress and site of failure in radial tension specimens (Skaggs et al., 1994a) and is believed to affect the meniscus’ free swelling strain (Andrews et al., 2015). Large transverse contractions in tensile tests are attributed to fiber architecture (Adeeb et al., 2004; Ban et al., 2019; Kiviranta et al., 2006; Reese et al., 2010). Loss of fiber continuity in a small specimen is likely to alter the meniscus’ mechanical response. The choice of specimen size and shape, therefore, is inevitably a compromise.

Regardless of which specimen geometry is used, the apparent Poisson ratio cannot be relied upon to predict outcomes in conditions that differ from the original test, and 3D constitutive models need to be developed for this purpose (Danso et al., 2015; LeRoux and Setton, 2002; Mononen et al., 2015, 2015; Párraga Quiroga et al., 2014; Spilker et al., 1992; Upton et al., 2006). Estimation of apparent Poisson ratios from planar uniaxial tension data for the meniscus did not produce consistent results, despite specimen dimensions differing by only 0.5 mm between studies (Goertzen et al., 1997; LeRoux and Setton, 2002). In tendon and ligament, which are also believed to be transversely isotropic, the apparent in-plane ν observed in uniaxial tension tests can range from approximately −3 to +6 (Cheng and Screen, 2007; Firminger and Edwards, 2021; Gatt et al., 2015; Hewitt et al., 2001; Luyckx et al., 2014; Lynch et al., 2003; Nagelli et al., 2022; Reese et al., 2010; Reese and Weiss, 2013; Swedberg et al., 2014). In some tissues, Poisson ratio, like the elastic modulus also appears to be strain-dependent (Chang et al., 1999; Edelsten et al., 2010; Elliott et al., 2002; Goertzen et al., 1997; LeRoux and Setton, 2002; Reese et al., 2010; Reese and Weiss, 2013; Swedberg et al., 2014; Woo et al., 1979), albeit with some findings to the contrary (Chang et al., 1999; Elliott et al., 2002; Huang et al., 2005). The present results support the general idea that circumferential tension tends to cause meniscus volume to decrease, but a 3D nonlinear constitutive model is required for precise, context-sensitive predictions. As it is not possible to fully characterize a 3D anisotropic material model using 2D data (Holzapfel and Ogden, 2009), the 3D data from the present work is also an important step towards developing a sound, evidence-based constitutive model for meniscus tissue.

### 4.3. Recovery

Physiologic loading of the knee meniscus is cyclic, so sustained fluid load support requires that the volume recovery rate during unloading be rapid enough that the meniscus can reset its volume and water content during the unloaded part of each cycle. In this study, the rate of volume change varied a great deal during both the loaded holds and unloaded recovery (Figure 8), which is expected. At the beginning of each loaded hold or unloaded recovery the state is far from equilibrium, so the rate of change is fast. As the equilibrium state is approached through volume loss or recovery, the rate of change approaches zero. Therefore, in cyclic loading the rates of volume loss and recovery will balance at some ε_v_ < 0, and the volume strain will oscillate about this strain level, effectively a natural set point. Based on the present results, this set point is expected to lie between ε_v_ = −0.03 ± 0.03 (volume strain at the start of each loaded hold, the time of fastest volume loss) and ε_v_ = −0.08 ± 0.04 (volume strain at the end of the each loaded hold, the time of slowest volume loss), because the rate of volume recovery immediately upon unloading lies between these loss rates (Figures 5, 7, and 8). This prediction assumes that recovery occurs by swelling alone, without any active recovery mechanism like those that have been proposed for articular cartilage (Moore and Burris, 2017; Voinier et al., 2022).

Previously published in vivo data is somewhat consistent with this set point prediction. Human meniscus volume approached an asymptote of ε_v_ ≈ −0.08 during a 20 km run (Kessler et al., 2006). This asymptote is most likely a steady state value, not an equilibrium value, as the equilibrium axial compression modulus for human meniscus is 50–100 kPa and contact pressures even for walking are 0.5–2 MPa (Chia and Hull, 2008; Gilbert et al., 2014; Walker and Erkman, 1975; Warnecke et al., 2020). At the very least, the present study produced similar volume strains as observed in vivo and is therefore observing phenomena with physiologic relevance.

Fast recovery of volume has also been observed in articular cartilage and the intervertebral disc. In indentation creep experiments on articular cartilage, the rate of recovery of indentation depth immediately after unloading was faster than the majority of the creep curve (Mow et al., 1989; Sweigart and Athanasiou, 2005a, 2005b). In creep indentation of human femoral head articular cartilage, recovery equilibrium was achieved 50% faster than creep equilibrium (Athanasiou et al., 1994). In a confined compression creep experiment, recovery from creep-induced compression took about the same time as the original loss (Zarek and Edwards, 1963). Considering the intervertebral disc, human subject height (which reflects changes in intervertebral disc height) recovered faster upon lying down than height was lost over the course of the day (Eklund and Corlett, 1984). In cadaveric experiments, cyclic loading of bovine discs and articular cartilage reaches steady state at ~ 10% compression (McCutchen, 1962; Schmidt et al., 2016), similar to our predictions for the meniscus above. Recovery may be described as slow when it is measured as the time to reach a free or fixed swelling equilibrium state (Supplemental Figure 2) (Bezci et al., 2020; Bezci and O‘Connell, 2018; O‘Connell et al., 2011; Schmidt et al., 2016), but for strains within the physiologic operating envelope volume recovery is often faster than its loss.

### 4.4. Swelling

Tissue tends to swell when stored unloaded in fluid (“free swelling”) (Andrews et al., 2015; Costi et al., 2002; Han et al., 2000; Han et al., 2012; Safa et al., 2017; Werbner et al., 2022, 2019; Żak and Pezowicz, 2016). However, repeated loading–recovery cycles in this study were on average sufficient to maintain the initial hydration level, with some specimens swelling and others shrinking over multiple cycles. Preconditioning appeared to reduce the impact of this variation by partially saturating the specimen’s tendency to swell or shrink prior to the test phases used for data analysis. Preconditioning with physiologic deformation/load and avoidance of free swelling, as done here, is a common recommendation and is generally successful at avoiding supraphysiologic swelling (Adams et al., 2000; Costi et al., 2021; Johannessen et al., 2006; Schmidt et al., 2016; Showalter et al., 2014).

### 4.5. Limitations

The major limitations of the present study are: (1) the ε_z_ and thus ε_v_ measurement rate was slow (sample period = 15–30 s), which prevented observation of dynamic (high strain rate) behavior, and (2) only a single strain level was tested. The slow sampling rate was due to using a DSLR camera, which at the time was necessary to achieve both high resolution and macro magnification. Better imaging options have since become commercially available, in principle allowing analysis of ~ 0.5 s ramps, matching gait loading (Chia and Hull, 2008; Uezaki et al., 1979). A single strain level was tested because manually labeling the z-edges and correcting inconsistencies between frames took 20–100 h per specimen (most of this variation was between workers). Cost-effective measurement of 3D strain with this approach depends on future development of automated analysis procedures.

### 4.6. Conclusion

Meniscus tissue loaded in circumferential tension was shown to lose volume immediately on loading and as the load is held, indicating fluid pressurization and exudation. Compression is already known to cause volume loss, so the combination of axial compression and circumferential tension that occurs in vivo should also cause volume loss. The meniscus is therefore expected to benefit from interstitial fluid pressurization and fluid load support, which provides lubrication and reduces load on the solid matrix, much like articular cartilage. Recovery of lost volume upon unloading from physiologically relevant volume strain (≈ −0.08) was rapid, faster than volume loss at physiologic volume strain, but slightly slower than the initial rate of volume loss upon loading from the zero strain initial condition. These results suggest that in cyclic loading volume loss and recovery will balance at moderate compressive strain, supporting maintenance of meniscus volume during sustained cyclic loading. Although the meniscus is believed to be transversely isotropic, transverse strain was found to significantly differ from through-thickness strain. Predictions of volume strain and thus fluid flow from planar (2D) data will therefore be very inaccurate, which is unfortunate because most prior measurements of meniscus deformation in tension are planar. It is essential to measure all three normal strain components in any work intended to investigate meniscus poroelasticity, and probably also in work with other poroelastic tissues.

## Supporting information

Plots of stress and strain vs. time for all specimens

## 5. Acknowledgments

Research reported in this publication was supported by (a) NIAMS of the National Institutes of Health under award number R21 AR070966, (b) NIGMS of the National Institutes of Health under award number U54 GM104941, and (c) the DCMR COBRE program under a grant from the National Institute of General Medical Sciences of the National Institutes of Health, NIH-NIGMS COBRE P20 GM139760.

## 1. Characteristic time results

Traditionally, the kinetics of a tissue’s viscoelastic response are quantified using exponential time constants (τ) measuring the time to achieve equilibrium, sometimes using a generalized Maxwell model [Bezci & O‘Connell 2018; Maritz … Barrera 2021; O‘Connell … Elliott 2011]. Exponential fits are useful if the data genuinely exhibits exponential decay and approximately reaches equilibrium. In the present experiment, strain components returned to approximately zero by the end of the unloaded holds, indicating equilibration at least for recovery. It is therefore potentially useful to report time constants for comparison to prior work. The decay was not always exponential, so an equivalent model-free metric was constructed: the time to achieve 63% of the total change, referred to as the “characteristic time” t_Δ63%_. If monoexponential decay truly describes the data the characteristic time t_Δ63%_ will be equal to the time constant τ of exponential decay. Otherwise, t_Δ63%_ can be interpreted the same way as τ; it simply avoids pathological results like τ ≈ 0 or τ → ∞ when the decay is not exponential.

The characteristic times of ε_v_ and ε_z_ were significantly longer during unloaded recovery (∼ 450 s) than during the loaded hold (∼ 170 s) (Figure 2); p < 0.005, stress relaxation and creep). The long characteristic time for recovery is in part attributable to recovery being held long enough to reach equilibrium, dε/dt ≈ 0 (see main text), whereas the loaded hold was not held for long enough to reach equilibrium. Therefore the characteristic times of loss and recovery cannot be compared on an equal basis, which is why rates of change were used in the main analysis.

**Figure 1:**
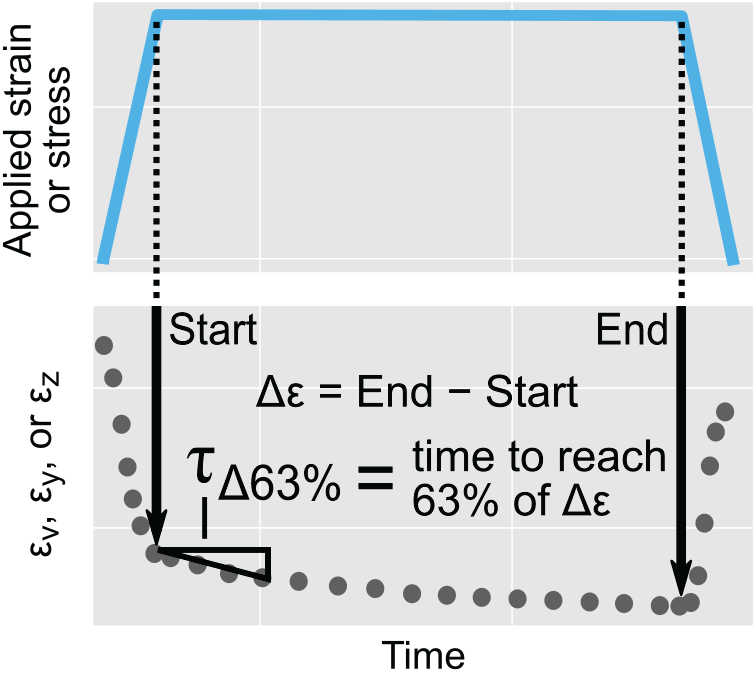
Schematic of calculation of the characteristic time τ_Δ63%_, defined as the time required for a strain or stress measurement to change by 63% of its total change during a loaded or unloaded hold.

**Figure 2:**
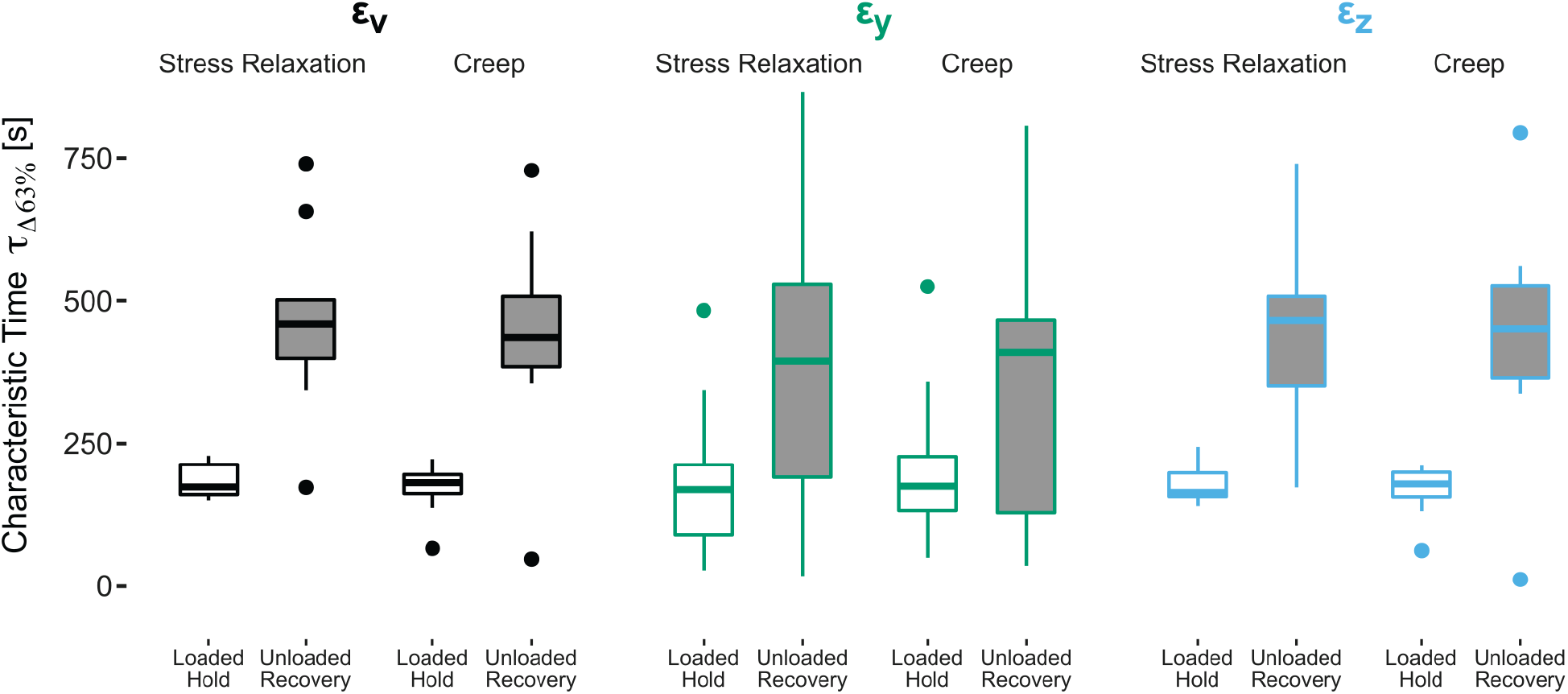
Characteristic time of change for each strain component during the loaded holds and unloaded recovery.

## 2. “Verification” stress relaxation phase results

In any test with repeated loading cycles, there is the potential for damage or other unrecoverable changes to accrue during the test, complicating comparison between cycles. In this work, the main stress relaxation and creep phases were compared and sometimes pooled, so it is important to verify that the specimen’s mechanical properties did not change during these phases. This verification was done by repeating the stress relaxation loading pattern at the very end of the test. The verification stress relaxation was within ∼ 0.1 MPa of the main stress relaxation (Figure 3), indicating that the main stress relaxation and creep phases produced negligible damage or other unrecoverable changes. The main stress relaxation and creep phases may therefore be compared on an equal basis.

**Figure 3:**
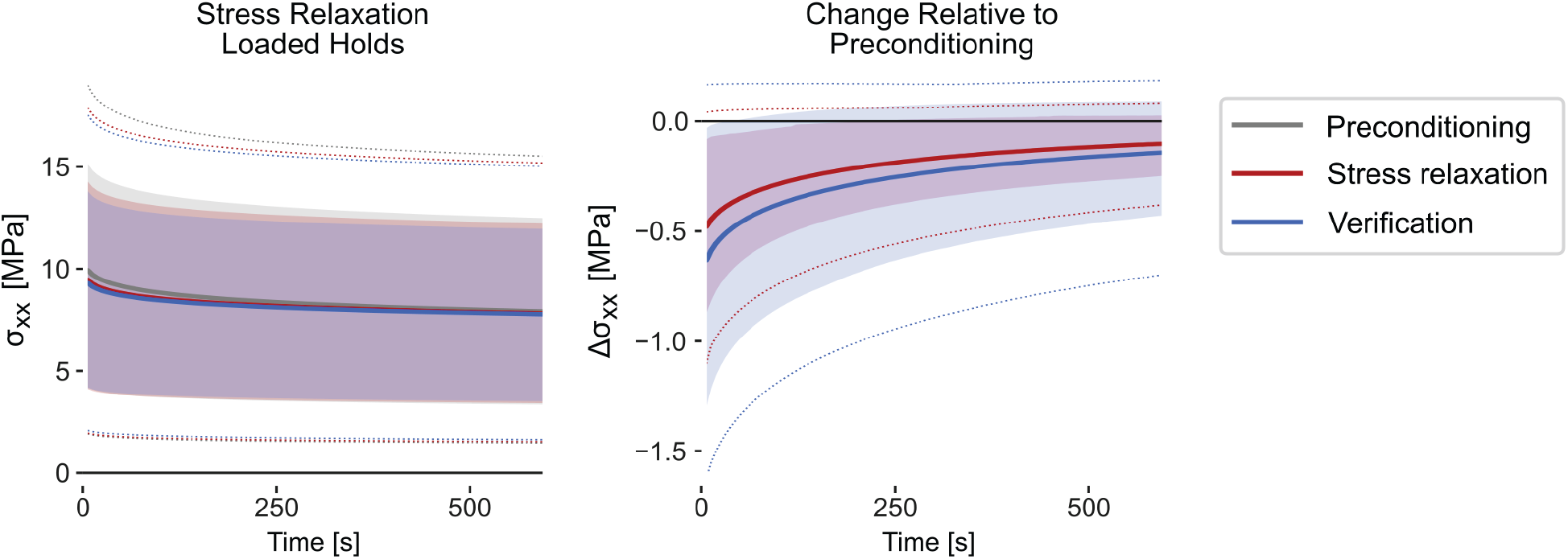
Comparison of all three stress relaxation-type loaded hold segments: preconditioning, the main stress relaxation phase, and the final verification phase. Solid line = mean across specimens. Shaded region = interquartile range. Dotted lines = min/max. The main stress relaxation and verification phases produced very similar stress values, slightly less than the preconditioning phase, indicating that the specimen accrued negligible damage or other unrecoverable changes during the main phases of the test protocol.

## Notes

### Competing Interest Statement

The authors have declared no competing interest.

